# Intraperitoneal oil application causes local inflammation with depletion of resident peritoneal macrophages

**DOI:** 10.1101/2020.07.15.203885

**Authors:** Elisenda Alsina-Sanchis, Ronja Mülfarth, Iris Moll, Carolin Mogler, Juan Rodriguez-Vita, Andreas Fischer

## Abstract

Oil is frequently used as a solvent to inject lipophilic substances into the peritoneum of laboratory animals. Although mineral oil causes chronic peritoneal inflammation, little is known whether other oils are better suited. Here we show that olive, peanut, corn or mineral oil causes xanthogranulomatous inflammation with depletion of resident peritoneal macrophages. However, there were striking differences in the severity of the inflammatory response. Peanut and mineral oil caused severe chronic inflammation with persistent neutrophil and monocyte recruitment, expansion of the vasculature and fibrosis. Corn and olive oil provoked no or only mild signs of chronic inflammation. Mechanistically, the vegetal oils were taken up by macrophages leading to foam cell formation and induction of cell death. Olive oil triggered caspase-3 cleavage and apoptosis, which facilitates the resolution of inflammation. Peanut oil and, to a lesser degree, corn oil triggered caspase-1 activation and macrophage pyroptosis, which impairs the resolution of inflammation. As such, intraperitoneal oil administration can interfere with the outcome of subsequent experiments. As a proof-of-principle, intraperitoneal peanut oil injection was compared to its oral delivery in a thioglycolate-induced peritonitis model. The chronic peritoneal inflammation due to peanut oil injection impeded the proper recruitment of macrophages and the resolution of inflammation in this peritonitis model. In summary, the data indicate that it is advisable to deliver lipophilic substances like tamoxifen by oral gavage instead of intraperitoneal injection.

## Introduction

Oil is frequently used as solvent in animal research. For instance, inducible gene recombination using the Cre-ERT2 - loxP system requires administration of tamoxifen which is usually dissolved in olive, peanut, corn or mineral oil. The oil solution is administered orally or by intraperitoneal injection (*i.p.*) (1, 2). Also, in a liver fibrosis model carbon tetrachloride (CCl_4_) is delivered by *i.p*. injection in oil, inhalation or oral gavage (3). Interestingly, *i.p*. injection generates stronger liver fibrosis when compared with the other two administration methods (4), raising the question whether CCl_4_ or its solvent act locally within the peritoneum. Indeed, *i.p.* injection of mineral oil causes chronic inflammation (5–9). Also, subcutaneous injection of olive oil can cause lipogranuloma, a granulomatous inflammatory soft tissue reaction (10).

Therefore, it can be assumed that any experimental immune cell analysis within the peritoneal cavity would be strongly affected by oil. It is surprising how little is known about the peritoneal immune cell reaction towards oil and comparative studies of different oils are missing to our knowledge.

Peritoneal inflammation can be divided into the initiation and resolution phase. Pathogens trigger infiltration of neutrophils, which phagocytose pathogens, clear apoptotic cells and recruit monocytes from the blood stream into the peritoneal fluid. Recruited monocytes eliminate dying neutrophils and differentiate into monocyte-derived macrophages (11). This is important, as the number of resident peritoneal macrophages, which are derived from embryonic progenitors and have self-renewal capacity (12), get strongly decreased as a result of the so-called “macrophage disappearance reaction” (13). As such, resident peritoneal CD11b^+^ macrophages, expressing high F4/80 levels (F4/80^hi^) get replaced by monocyte-derived CD11b^+^ macrophages, expressing low F4/80 levels (F4/80^low^) on the membrane (12, 14, 15). Subsequently, monocyte-derived macrophages increase surface expression of F4/80 from a low to an intermediate level (F4/80^int^) to initiate the resolution phase (16).

The switch from inflammatory to resolving macrophages is triggered by phagocytosis of apoptotic cells. Deficiency in this phagocytic process leads to chronic inflammation (17). For instance in atherosclerotic plaques, macrophages take up excessive amounts of lipids and become foam cells, which cannot initiate the resolution phase, perpetuating further neutrophil and monocytes infiltration (18).

The aim of this study was to analyze how the most commonly used oils in animal research affect the myeloid cells within the peritoneum and whether this would diminish their capability to resolve the peritoneal inflammation.

## Materials and methods

### Animal models

The study was approved by institutional and regional animal research committees. All animal procedures were in accordance with institutional guidelines and performed according to the guidelines of the local institution and the local government. Female C57BL/6 mice were group‐housed under specific pathogen‐free barrier conditions.

Administration of peanut (P2144, Sigma-Aldrich, St. Louis, USA), corn (C8267, Sigma-Aldrich, St. Louis, USA), olive (88631, Carl Roth, Germany), mineral oil (HP50.2, Carl Roth, Germany) or 0,9% sterile NaCl (Braun, Germany) in 8 to 12-week-old randomized mice was performed by daily *i.p*. injection of 100 μl for 5 consecutive days or by oral gavage of peanut oil once with 100 μl. After three weeks mice were euthanized. For peritoneal lavage, 5 ml of cold PBS (Gibco/Thermo Fisher Scientific, NY, USA) was injected *i.p*. after a careful massage to mobilize cells, peritoneal fluid was collected. Cells were isolated by centrifugation (5 min, 200 g) and suspended in 1 ml of PBS.

8 to 12-week-old randomized mice were euthanized and administrated with peanut, olive, corn and mineral oil. After 5 minutes the peritoneal lavage was collected.

Three weeks after oil treatment, mice were *i.p*. injected with thioglycolate (2 mg in 1 ml H_2_O; B2551, Sigma Aldrich, St. Louis, USA). After 24 or 72 hours mice were sacrificed and peritoneal lavage collected. All groups were randomized.

### Immunofluorescence and tissue histology

Histological analysis was performed on formalin‐fixed paraffin‐embedded sections (3 μm). Sections were deparaffinized and rehydrated. For hematoxylin-eosin (H&E) and Sirius Red (Dianova, Germany) staining, sections were processed according to standard protocols. For myeloid cell staining, antigen retrieval at pH 6 with citrate buffer and the primary antibody rabbit anti-mouse CD11b (1:200) (ab133357, Abcam, Cambridge, MA, USA) and antigen retrieval with 1:20 proteinase K/TE buffer and rat anti-mouse F4/80 (1:100) (T-2006, Dianova, Germany) incubated at 4°C overnight. After washing, sections were incubated with secondary antibodies coupled with HRP (1:200) (DAKO, Agilent Technologies, Santa Clara, CA, USA) for one hour at room temperature. For immunofluorescence staining, antigen retrieval at pH 9 was performed using citrate buffer and sections were incubated with the primary antibody rabbit anti‐mouse CD31 (1:50) (ab28364, Abcam, Cambridge, MA, USA) at 4°C overnight. After washing, sections were incubated with secondary antibody (1:200) goat anti‐rabbit Alexa Fluor-647 (A21245, Life Technologies/Thermo Fisher Scientific, NY, USA) for 1 hour at room temperature. H&E images were obtained with slide scanner (Zeiss Axio Sacn.Z1, Carl Zeiss, Germany). CD11b images were obtained with widefield microscope (Zeiss Axioplan, Carl Zeiss, Germany). All images were processed with ZENblue software (Carl Zeiss, Germany). Immunofluorescence was imaged at the confocal (LSM 700, Carl Zeiss, Germany) microscope with ZENblack software (Carl Zeiss, Germany). Sections of seven Z-stacks per omentum and mesentery and three random fluorescence images per slide were taken. Numbers of CD31 positive vessels per view field and lipid droplet size from H&E images were counted with ImageJ software (NIH, Bethesda, MD, USA).

### Oil Red O staining

Peritoneal lavage was plated into one well of a 6-well plate on top of coverslips and incubated for 30 min with Dulbecco’s modified Eagle’s medium (DMEM) (Gibco/ Thermo Fisher Scientific, NY, USA). Afterwards non-adherent cells were removed by careful washing three times with PBS. J774A.1 cells cultured in DMEM with 10% fetal calf serum (Biochrom, UK) were seeded into 12-well plates on coverslips and treated with 100 μl oil in 1 ml medium for four hours. Cells on coverslips were stained with Oil Red O (O0625, Sigma-Aldrich) following the protocol published elsewhere (19) and counterstained with hematoxylin. Images were obtained with widefield microscope (Zeiss Axioplan Carl Zeiss, Germany).

### Flow cytometry

Cells obtained from peritoneal lavage were washed and erythrocytes lysed with ACK lysis buffer (Gibco/Thermo Fisher Scientific, NY, USA). Cells were suspended at approximately 10^6^ cells/ml in PBS with 2% FCS. Cell suspensions were incubated with the different fluorophore-coupled primary antibodies for 20 minutes on ice. These antibodies were used: CD45 (552848), CD11b (552850), CD19 (560375), Ly6G (560600), Ly6C (560594) and F4/80-like (564227) all from BD Biosciences (Bedford, MA, USA), CD3 (100203) and F4/80 (123128) from BioLegend (St. Diego, CA, USA) and Tim4 (12-5866-82, Life Technologies/Thermo Fisher Scientific, NY, USA). Concentration of the different antibodies was determined by titration. Flow cytometer results in percentage were extrapolated to the total amount of cells obtained from the previous cell counting.

### Western Blot analysis

Cell lysates were separated by SDS-PAGE and proteins blotted on nitrocellulose membranes. Membranes were blocked with 5% skim milk in TBS with 1% Tween-20. The following primary antibodies were used: CD36 (ab124515), VCP (ab11433) from Abcam (Cambridge, MA, USA), ABCG1 (NB400-132SS, Novus Biologicals, CO, USA), Cleaved-Caspase 3 (Asp175; 9664S), Arginase-1 (D4E3M™; 93668S) from Cell Signaling (Danvers, MA, USA) and Caspase 1 (14F468; sc-56036, Santa Cruz Technologies, Dallas, TX, USA). Primary antibodies were incubated overnight at 4°C and appropriate HRP-conjugated secondary antibodies (DAKO, Agilent Technologies, Santa Clara, CA, USA) for 1 hour at room temperature. Chemiluminescence was detected by Pierce ECL Western Blotting Substrate (Thermo Fisher Scientific, NY, USA) and ChemiDoc imaging system (Biorad, Hercules, CA, USA) and quantified with Image Lab 3.0 software (Biorad, Hercules, CA, USA).

### Quantitative PCR

RNA was isolated using the innuPREP RNA Mini kit (Analytik Jena, Germany). cDNA was synthesized with the High‐Capacity cDNA Reverse Transcription Kit (Applied Biosystems). The cDNA was applied to qPCR using the POWER SYBR Green Master Mix (Applied Biosystems). Fold changes were assessed by 2^−ΔΔ*C*t^ method and normalized with the *CPH* gene. The following primers were used for qPCR: CD36 forward GCAAAACGACTGCAGGTCAA and reverse GGCCATCTCTACCATGCCAA, ABCG1 forward CTTTCCTACTCTGTACCCGAGG and reverse CGGGGCATTCCATTGATAAGG, IL10 forward GCATGGCCCAGAAATCAAGG and reverse GAGAAATCGATGACAGCGCC and CPH forward ATGGTCAACCCCACCGTG and reverse TTCTTGCTGTCTTTGGAACTTTGTC.

### Cell death detection

J774A.1 cells were plated at 5×10^5^ cells per well into a 12 well-plate with 100 μl of the different oils in 1 ml medium and incubated for four hours. Afterwards, supernatant and attached cells were collected and stained with Annexin V-FITC (640905, BioLegend, St. Diego, CA, USA) and PI (Cayman Chemical, USA) and incubated for 15 minutes on ice. After washing, cells were immediately analyzed by flow cytometry.

An apoptosis/necrosis immunofluorescence assay kit (ab176749, Abcam, Cambridge, MA, USA,) was used for detection of necrosis or apoptosis cell death. J774A.1 cells were plated at 5×10^5^ cells/well into a 24 well-plate on top of a coverslip. To each well, 50 μl of oil was added in a final volume of 1 ml medium and incubated for two hours. Afterwards, the staining was performed following the manufacturer’s protocol. Three fluorescence images of each channel at fixed positions of each triplicate were collected at the wide-field Cell Observer microscope (Carl Zeiss, Germany) with ZENblue software (Carl Zeiss, Germany). FIJI software was employed for the quantification of positive cells of each channel per field.

For determination of lactate dehydrogenase activity in the cell supernatant, J774A.1 cells were plated at 5×10^4^ cells/well into a 96 well-plate with 100 μl of medium containing 10 μl oil in triplicates and incubated for 2 hours. Oleic acid (O1008, Sigma-Aldrich, St. Louis, USA), diluted in absolute ETOH was added to the medium or mixed with 5 μl peanut oil when indicated. Then, levels of LDH were detected using the LDH-Cytotoxicity Assay Kit (Ab65393, Abcam, Cambridge, MA, USA) following the manufacturer’s protocol.

### Statistical analysis

GraphPad Prism 8 (GraphPad Software, San Diego, CA, USA) was used to generate graphs and for statistical analysis. Statistical significance was calculated using one-way or two-way ANOVA as indicated in the figure legends. Data sets are presented as mean ± SD. *P* < 0.05 was considered as significant.

## Results

### Macroscopic changes upon intraperitoneal oil injection

Peanut, olive, corn and mineral oil were injected into the peritoneum (*i.p.*) of adult mice for five consecutive days. This mimics a typical protocol for delivering tamoxifen to induce gene recombination in transgenic mice expressing Cre^ERT2^ recombinase (2). Analysis was done three weeks later. As controls, untreated mice and mice treated with peanut oil by oral gavage were used (**Figure 1A**).

**Figure 1.**
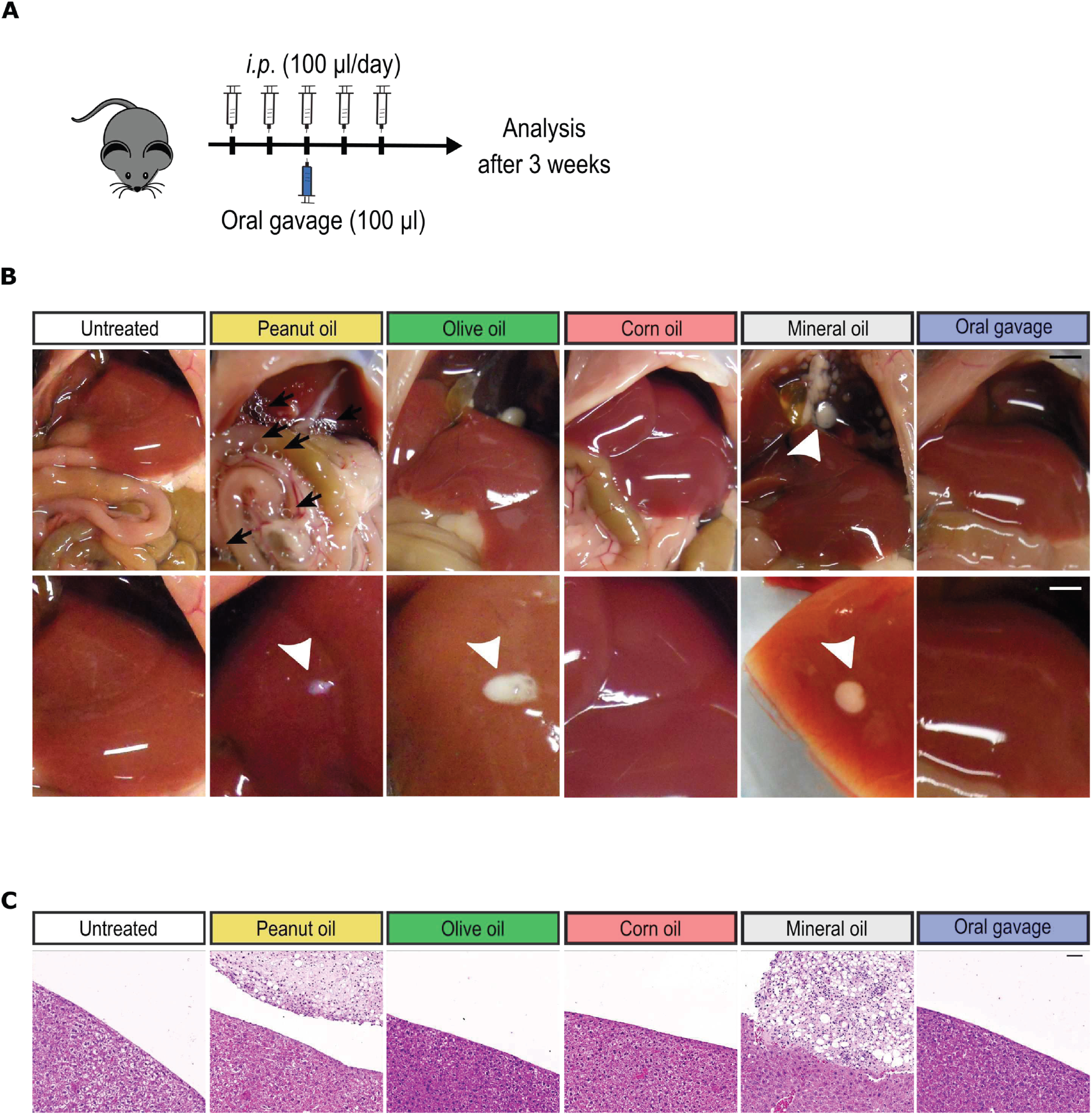
Macroscopic changes in mice three weeks after *i.p.* oil injection. **A**. Schematic illustration of the *i.p.* or oral oil administration protocol. **B.** Representative images of the peritoneum. Black arrows indicate visible lipid droplets. White arrowheads mark nodules on the surface of organs. Scale bar, 3 mm. **C**. Representative microscopic images of liver sections stained with H&E. Xanthogranuloma on the liver surface in mice treated with peanut and mineral oil. Scale bar, 50μm.

In contrast to untreated mice or those receiving oil by oral gavage, the *i.p.* injected mice showed macroscopically visible alterations in the peritoneal cavity. Peanut oil was still visible as oil droplets (**Figure 1B**), whereas this was not the case for the other oils. White nodules, in the size of <3 mm, were visible on the surface of liver, diaphragm or colon in mice receiving peanut, olive and mineral oil *i.p.* but not corn oil. The nodules formed due to olive oil treatment were only loosely attached to the organ surfaces, whereas the nodules in mice that were *i.p.* injected with peanut or mineral oil were firmly attached to the liver surface (**Figure 1B**).

### Xanthogranulomatous inflammation in the peritoneum upon oil injection

Histological analysis revealed no pathological changes in liver (**Figure 1C**) and spleen (**Supplementary Figure 1A**) of mice that received *i.p.* injection of oil. The nodules that were firmly attached to the liver surface in mice treated with peanut or mineral oil could be classified as xanthogranulomas with foamy macrophages and mixed inflammatory background (**Figure 1C**).

Next, we examined the greater omentum. The greater omentum is an organ that filters excessive fluid from the abdominal cavity, senses microorganisms or damaged cells, initiates immune responses and supports repair of damaged organs (20). Omenta of mice treated with oral gavage were indistinguishable from those of untreated mice. However, omenta of mice that received *i.p.* oil injections showed a remarkable change in morphology. The omenta were swollen, darker and had enlarged blood vessels, particularly in mice treated with peanut and mineral oil (**Figure 2A**). Histological analysis revealed that lipid droplet size in adipocytes was reduced in mice treated *i.p.* with any of the different oils. Again, the changes were most pronounced in mice injected with peanut and mineral oil (**Figures 2B and 2F**).

**Figure 2.**
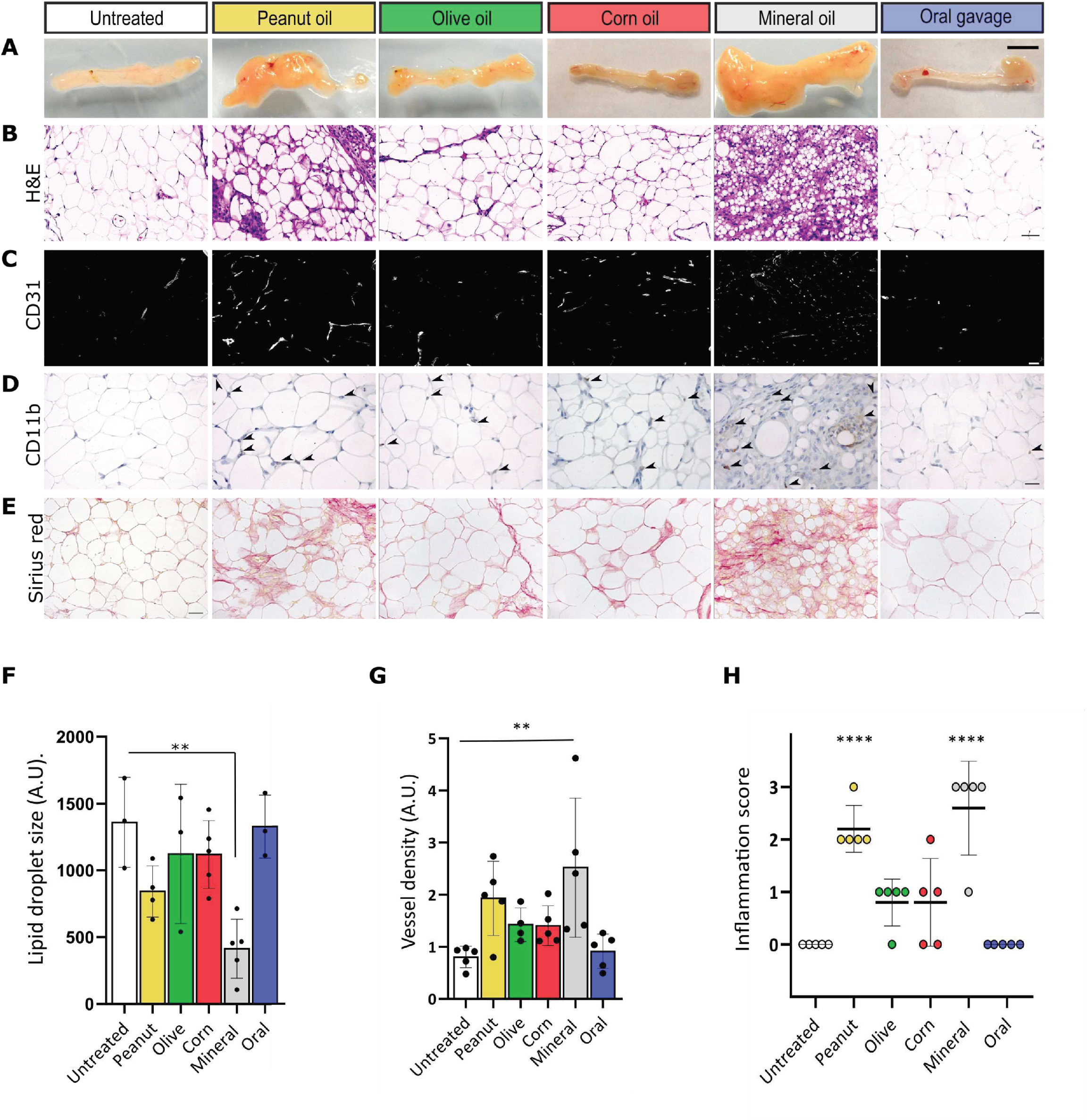
Xanthogranulomatous inflammation in the omentum upon oil injection. **A**. Representative images of omenta from untreated mice and such that were treated with oil by *i.p.* injection or oral gavage. Analysis was performed three weeks after treatment. **B.** Representative confocal images of mesentery sections stained with H&E. Scale bar, 50 μm. **C**. Immunofluorescence microscopy to detect CD31^+^ endothelial cells (white). Scale bar 20 μm. **D**. CD11b^+^ myeloid cells. Scale bar, 100 μm. **E** Sirius Red staining to detect fibrosis. Scale bar, 100 μm. **F**. Lipid droplet size quantification from H&E images. Untreated micen=3, peanut oil n=4, olive oil n=3, corn oil n=5, mineral oil n=5 and oral gavage n=3. Bar graphs show mean ± SD, *, p<0.05 (one way ANOVA). **G**. Quantification of microvessel density. Olive oil n=4 and all other groups n=5. Bar graphs show mean ± SD, *, p<0.05 (one way ANOVA). **H**. Xanthogranulomatous inflammation score. 0: no inflammation to 3: very strong inflammation. All data represent n=5 mice, bar graphs show mean ± SD, *, p<0.05 (one way ANOVA).

Immunohistochemical analysis of CD31-positive endothelial cells revealed an increase in vessel density in the omenta after oil injection. It was strongly increased in the case of peanut and mineral oil but mild in olive and corn oil-treated mice (**Figure 2C and 2G**). In addition, higher numbers of CD11b^+^ myeloid cells were present in all four oil-treated mice, but again, peanut and mineral oil-treated mice had highest infiltration rates (**Figure 2D**).

So far, the described changes are indicative of peritoneal inflammation upon local oil injection. Prolonged inflammation may impede tissue healing resulting in organ fibrosis. Indeed, Sirius Red staining revealed an increase in collagen deposition, a typical sign of fibrosis, in the greater omentum of *i.p.* oil-injected mice. Such fibrotic changes were in particular observed in mice treated with peanut and mineral oil (**Figures 2E**).

Histopathological scoring of the inflammation grade in the greater omentum by H&E staining was based on the granularity of the tissue, the presence of foamy macrophages or other inflammatory cells, multinucleated giant cells, fibrosis or necrosis with a score from 0 (no inflammation) to 3 (severe chronic inflammation). This showed that *i.p.* injection of all four oils causes xanthogranulomatous inflammation of the omentum with highest scores for mineral and peanut oil. (**Figure 2H**).

Similar data were obtained during the analysis of mesentery. The almost transparent membrane became opaque in mice treated *i.p.* with oil. The strongest changes were observed in mice treated with peanut and mineral oil (**Figure 3A**). Lipid droplets in adipocytes of mesentery from mineral oil-treated mice were much smaller compared to controls. Such changes were also observed but to a lesser extent in mice injected *i.p.* with peanut oil, whereas the effects of olive and corn oil were mild (**Figures 3B and 3F**). The number of blood vessels were increased in peanut and mineral oil-treated mice. We detected increased numbers of CD11b^+^ myeloid cells in the mesentery of olive, corn, peanut and to the maximum extent in mineral oil-treated mice (**Figure 3D**). There was mild fibrosis in the mesentery of mice treated with peanut oil and severe fibrosis in mice that had received mineral oil (**Figure 3E**). Histopathological scoring of the inflammation grade revealed that peanut and mineral oil, but not olive and corn oil, generated xanthogranulomatous inflammation (**Figure 3H**).

**Figure 3.**
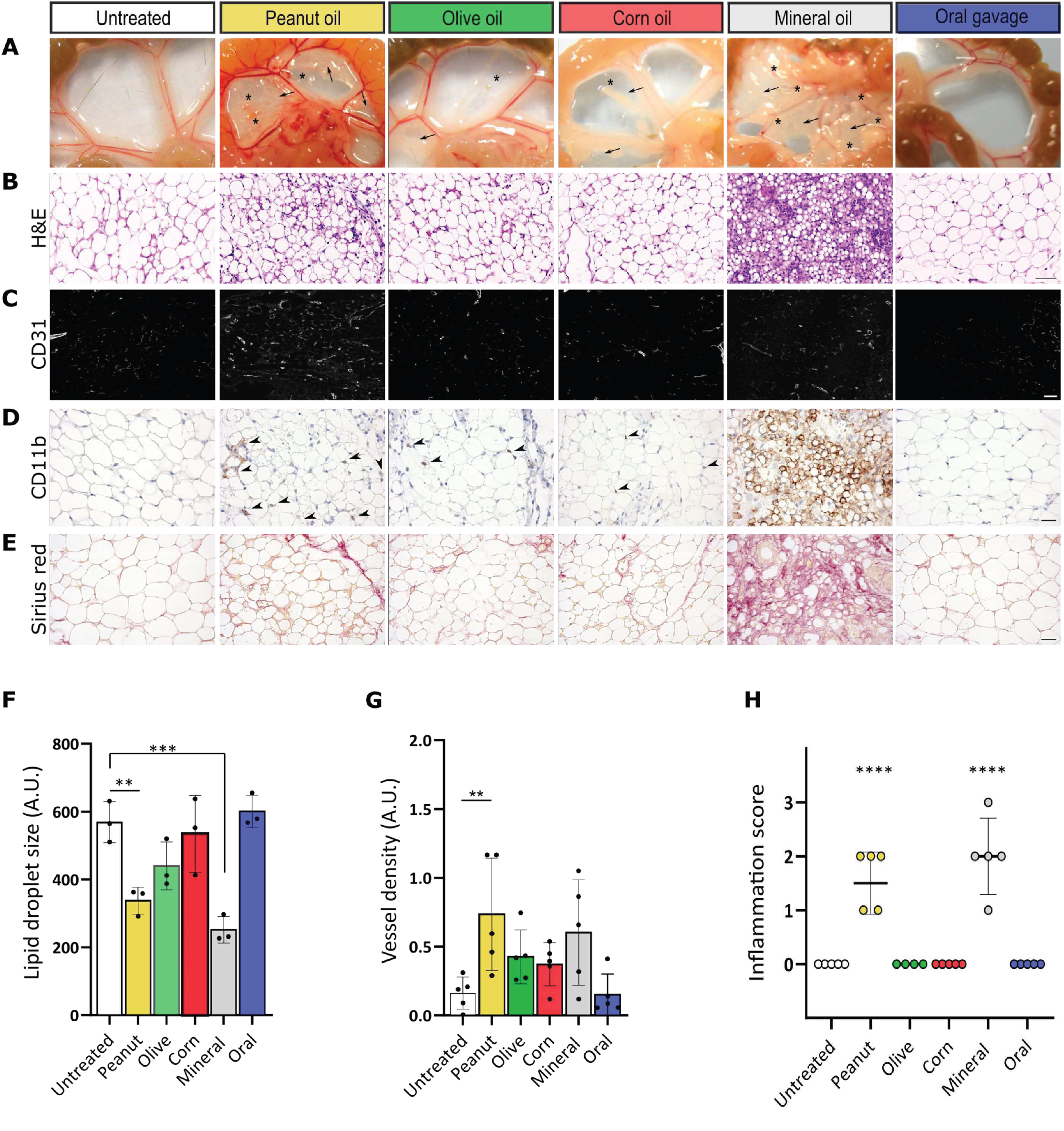
Xanthogranulomatous inflammation in the mesentery upon peanut and mineral oil injection. **A**. Representative images of mesentery from untreated mice and such that were treated with oil by *i.p.* injection or oral gavage. Analysis was performed three weeks after treatment. **B.** Representative confocal images of mesentery sections stained with H&E. Scale bar, 50 μm. **C**. Immunofluorescence microscopy to detect CD31^+^ endothelial cells (white). Scale bar, 20 μm **D**. CD11b^+^ myeloid cells. Scale bar, 100 μm **E** Sirius Red staining to detect fibrosis. Scale bar, 100 μm **F**. Lipid droplet size quantification from H&E images. n=3 mice. Bar graph shows mean ± SD, *, p<0.05 (one way ANOVA). **G**. Quantification of microvessel density. n=5 mice. Bar graph shows mean ± SD, *, p<0.05 (one way ANOVA). **H**. Xanthogranulomatous inflammation score. 0: no inflammation to 3: very strong inflammation. Olive oil n=4 and all other groups n=5. Bar graph shows mean ± SD, *, p<0.05 (one way ANOVA).

In summary, the histopathological analysis revealed that mineral oil and peanut oil induce a strong xanthogranulomatous inflammatory response in the peritoneum. Olive and corn oils also induce inflammation, but to a much lesser degree.

### Intraperitoneal oil injection causes myeloid cell infiltration into the peritoneum

To further analyze the immune response, peritoneal lavage was obtained three weeks after the *i.p.* injection. Flow cytometry revealed that the total cell number in the peritoneal lavage was significantly increased in mice treated with either peanut or mineral oil compared to untreated animals (**Figure 4A**). The vast majority of the cell population were myeloid cells (CD45^+^CD19^−^CD11b^+^). Peanut and mineral oil increased the total number of myeloid cells, whereas olive and corn oil had no significant effect (**Figure 4B and Supplementary Figure 2A**).

**Figure 4.**
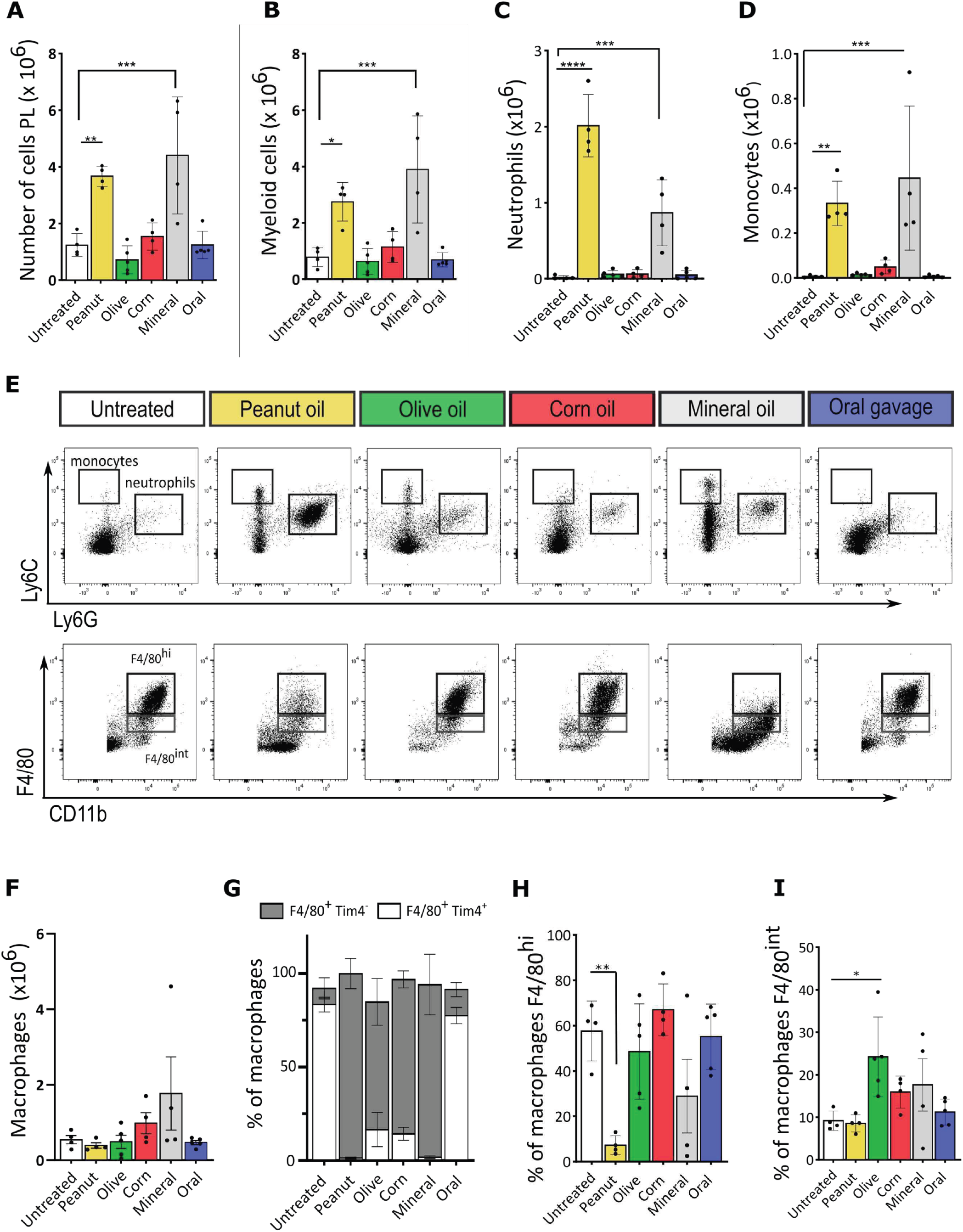
Resident peritoneal macrophage depletion and persistent monocyte and neutrophil infiltration upon peanut and mineral oil injection. Peritoneal lavage (PL) was obtained three weeks after treatment with oil. **A**. Total cell number in peritoneal lavage **B**. Myeloid cells (CD45^+^CD11b^+^) in peritoneal lavage. **C**. Neutrophils (CD45^+^CD11b^+^Ly6G^+^Ly6C^int^) in peritoneal lavage. **D**. Monocytes (CD45^+^CD11b^+^Ly6G^−^Ly6C^+^) in peritoneal lavage. **F**. Macrophages (CD45^+^CD11b^+^F4/80^+^) in peritoneal lavage. **E**. Representative blots of flow cytometry analysis of monocytes, neutrophils and macrophages. **G.** Percentage of CD45^+^CD11b^+^F4/80^+^Tim4^+^ and CD45^+^CD11b^+^F4/80^+^Tim4^−^ macrophages. **H.** Percentage of CD45^+^CD11b^+^F4/80^hi^ macrophages. **I.** Percentage of CD45^+^CD11b^+^F4/80^int^ macrophages. All data represent n=4 mice for untreated, peanut, corn and mineral groups and n=5 for olive and oral gavage groups. Bar graphs show mean ± SD, *, p<0.05 (one way ANOVA).

We also observed that oil injection led to a decrease in the number of B (CD45^+^CD19^+^CD3^−^) and T lymphocytes (CD45^+^CD19^−^CD3^+^) (**Supplementary Figure 2B**). This was expected as during peritoneal inflammation lymphocytes migrate from the peritoneal fluid into the greater omentum (21, 22).

### Peanut and mineral oil increases neutrophil and monocyte recruitment

A more detailed evaluation of the myeloid population revealed that peanut and mineral oil injection strongly increased the presence of neutrophils (CD45^+^CD11b^+^Ly6G^+^Ly6C^int^) and recruited monocytes (CD45^+^CD11b^+^Ly6G^−^Ly6C^+^) in the peritoneal fluid. Neutrophils and infiltrated monocytes were almost absent in peritoneal lavage derived from untreated mice or mice treated with olive oil, corn oil or oral gavage (**Figures 4C, D and E**).

Importantly, the injection itself did not cause such alterations. Injection of 0.9% NaCl did not lead to macroscopic or histological changes, nor to significant changes in total number of cells in peritoneal fluid or changes within the myeloid cell compartment (**Supplementary Figure 3A-G**).

### Oil injection leads to a severe reduction of resident peritoneal macrophages

During inflammation, neutrophils are the first cells being recruited to clear apoptotic cells or eliminate pathogens. Afterwards, monocytes reach the inflamed zone to eliminate dying neutrophils and to differentiate into macrophages. The latter is in particular essential when resident macrophages are eradicated. Therefore, we next examined the macrophage population within the peritoneum. The total macrophage (CD45^+^CD11b^+^F4/80^+^) cell number was not significantly changed in peritoneal lavage of mice treated *i.p.* with oil compared to the untreated mice or to those which received oil by oral gavage (**Figure 4F**).

We further characterized the F4/80 population by analyzing the amount of Tim4 on the cell surface as Tim4 can be employed as a marker to differentiate long-term (F4/80^+^Tim4^+^) from newly recruited (F4/80^+^Tim4^−^) resident macrophages (23). This revealed that all four oils led to a dramatic decrease in long-term resident (F4/80^+^Tim4^+^) macrophages and a replacement by recently recruited (F4/80^+^Tim4^−^) macrophages (**Figure 4G**).

The level of F4/80 on the macrophage cell membrane varies depending on the differentiation stage (15). In this regard, resident macrophages express high levels of F4/80, while newly recruited monocyte-derived macrophages express low to intermediate levels (F4/80^int^). Peanut oil injection led to a strong decrease in resident F4/80^hi^ macrophage numbers (**Figure 4H**). Mineral oil showed a similar but not significant trend, whereas olive oil and corn oil did not alter the proportion of F4/80^hi^ macrophages.

The full resolution of inflammation is carried out by F4/80^int^ macrophages (16). Analysis of this cell population revealed that only the *i.p.* injection of olive oil led to a significant increase in F4/80^int^ macrophages at this time point (**Figure 4I**).

Collectively, the data imply that injection of any oil into the peritoneum triggers an inflammatory response in which resident macrophages get replaced by monocyte-derived ones. However, the resolution of inflammation depends on the type of oil, with olive oil and corn oil (to a lesser extent) showing signs of resolution.

### Oil injection induces foam cell formation

Monocytes and macrophages can take up excessive amounts of lipids (24). Therefore, we examined lipid uptake in macrophages upon oil injection. Peritoneal lavage was performed three weeks after *i.p.* oil injection. Adherent peritoneal macrophages were stained with Oil Red O, which marks lipids and neutral triglycerides. Macrophages derived from the control mice were not stained by Oil Red O while peritoneal macrophages derived from mice injected *i.p.* with peanut oil contained multiple large lipid droplets. Fewer amounts were detected in the macrophages from the olive oil and corn oil group, whereas the macrophages of the mineral oil group contained almost no detectable lipid droplets (**Figure 5A**).

**Figure 5.**
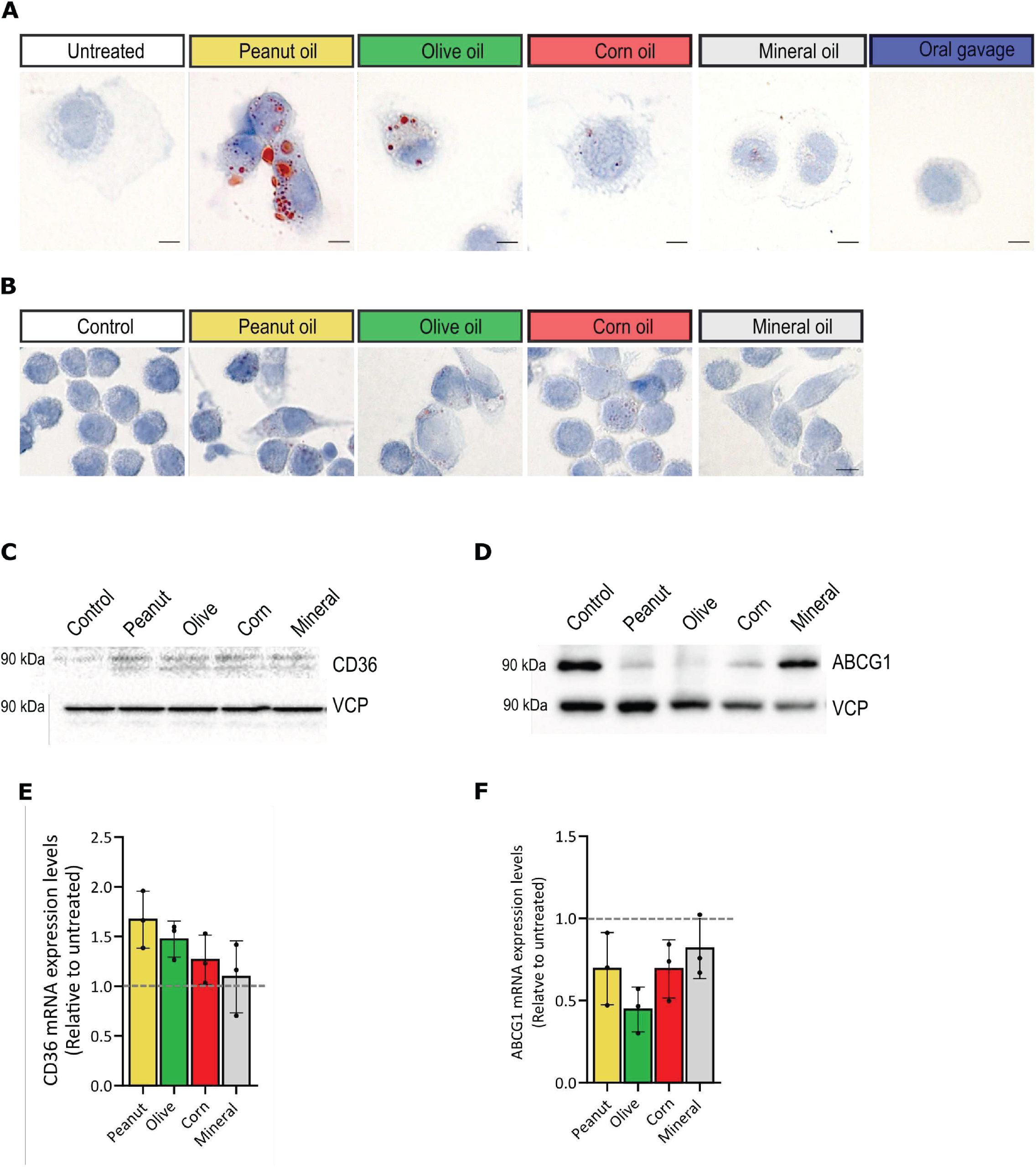
Macrophage foam cell formation upon contact with vegetal oil. **A**. Isolation of peritoneal macrophages from untreated mice and three weeks after peanut, olive, corn, mineral oil intraperitoneal injection. Cells from peritoneal lavage were plated for 30 minutes and stained with Oil Red O. Scale bar, 5 μm. **B.** J774A.1 macrophages after four hours in contact with the different oils. Representative images of Oil Red O staining. Scale bar, 50 μm. **C**. Representative Western blot of CD36 expression in J774A.1 macrophages after four hours in contact with the different oils. **D.** Representative Western blot of ABCG1 expression in J774A.1 macrophages after four hours in contact with the different oils. **E.** Quantification of CD36 and ABCG1 mRNA expression levels expression in J774A.1 macrophages after four hours in contact with the different oils. Fold change in comparison to untreated cells. All data from n=3 biological replicates, bar graphs represent mean ± SD, *, p<0.05 (one way ANOVA).

To further evaluate this, we tested lipid uptake in the J774A.1 macrophage cell line. J774A.1 macrophages took up lipids when in contact with olive, corn and peanut oil, but not mineral oil, suggesting that the effect of this oil is independent of the cellular lipid uptake.

In atherosclerotic plaques, monocyte-derived macrophages endocytose lipids such as oxidized LDL and become foam cells. During this transition, an upregulation of the fatty acid translocase (CD36) expression and downregulation of the cholesterol transporter ABCG1 is evident (25). J774A.1 macrophages showed the same changes in gene expression when cultured for four hours in the presence of vegetal oils (**Figure 5C-F**).

In summary, these results indicate that peanut, olive and corn, but not mineral oil *i.p.* injection leads to foam cell formation.

### Peritoneal macrophage cell death after exposure to different oils

Lipoprotein uptake can cause macrophage cell death (26). Therefore, we evaluated whether peritoneal macrophage cell death is induced by the four different oils. Mice were injected once *i.p.* with 100 μl oil and five minutes later peritoneal cells were harvested and subjected to flow cytometry (**Figure 6A**). This revealed that compared to untreated mice there was approximately a 50% decrease in CD11b^+^F4/80^+^ macrophages in the peritoneum of mice that received any of the oils (**Figure 6B**). There was a 5-10% decrease in the fraction of live cells (Annexin V^−^, PI^−^) and an equivalent increase of necrotic (PI^+^), apoptotic (Annexin V^+^) and Annexin V^+^PI^+^ cells in the peritoneal lavage of mice that received peanut or olive oil (**Figure 6C**). The Annexin V^+^PI^+^ double-positive population can be the result of both, apoptosis and necrosis (27). The increase in cell death was milder in the presence of corn oil, whereas in mineral oil injected mice cell death was not different as compared to untreated mice, suggesting that the mechanism for mineral oil-induced injury is different (**Figure 6C**).

**Figure 6.**
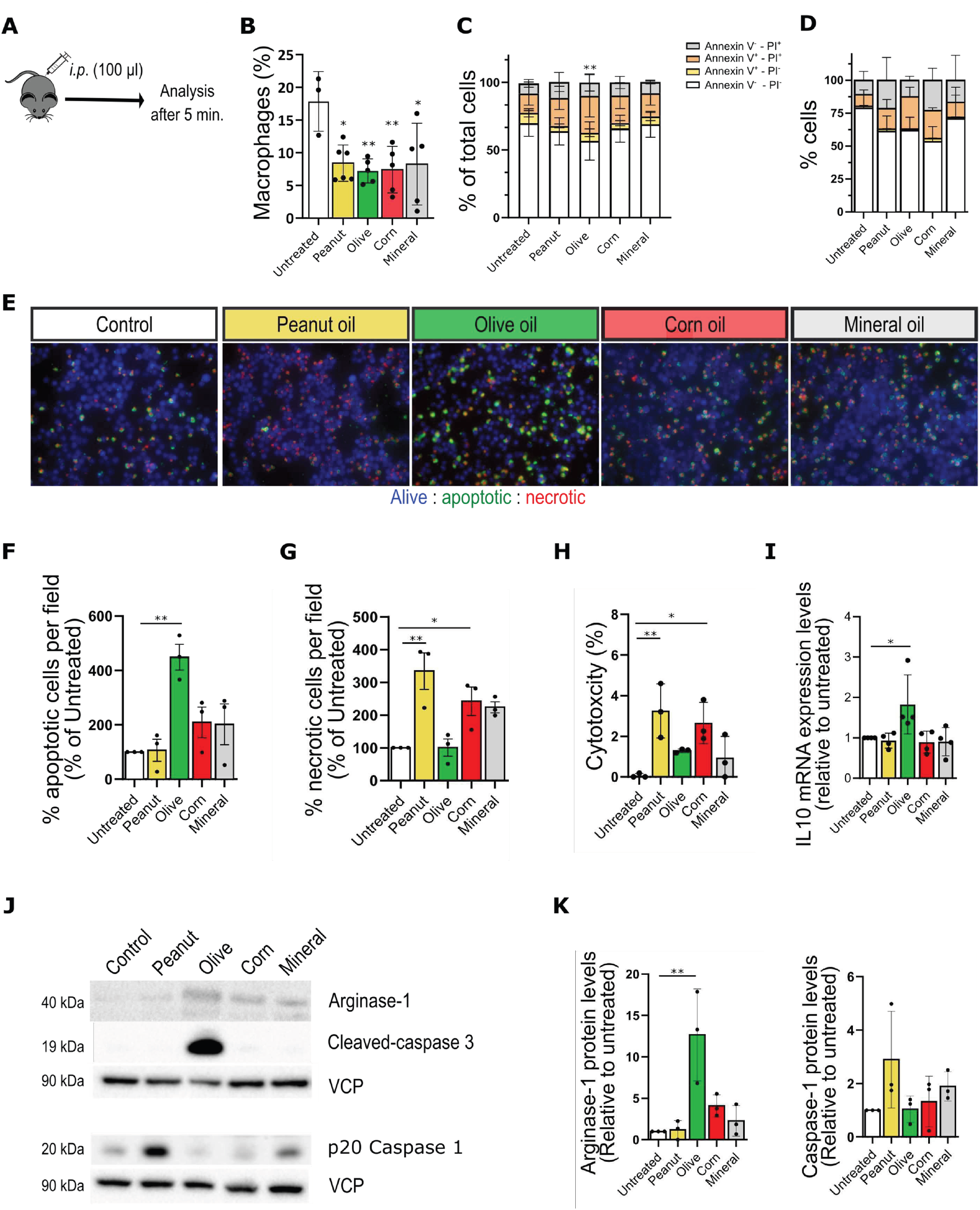
Macrophage cell death upon exposure to vegetal oil. **A**. Schematic illustration of *i.p.* oil administration. Peritoneal lavage was obtained five minutes later. **B**. Percentage of macrophages (CD45^+^CD11b^+^F4/80^+^). Untreated mice n=3, peanut oil n=6 mice and all other groups n=5 mice, mean ± SD, *, p<0.05 (one way ANOVA). **C**. Percentage of live (Annexin V^−^ PI^−^), apoptotic (Annexin V^+^PI^−^), necrotic (Annexin V^−^PI^+^) or double positive (Annexin V^+^PI^+^) cells from total number of cells in peritoneal lavage. Untreated mice n=6, peanut and olive oil n=9 mice, corn and mineral oil n=10 mice, bar graphs represent mean ± SD, *, p<0.05 (2-way ANOVA). **D.** J774A.1 macrophage cell line untreated or incubated with different oils for four hours and analysis of the percentage of live (Annexin V^−^PI^−^), apoptotic (Annexin V^+^PI^−^), necrotic (Annexin V^−^PI^+^) or double positive (Annexin V^+^PI^+^) cells. n=3 biological replicates, bar graphs represent mean ± SD, *, p<0.05 (2-way ANOVA). **E.** J774A.1 macrophages untreated or incubated with different oils for two hours. Quantification positive cells per field in comparison to untreated, blue (alive cells), green (apoptotic) and red (necrotic). n=3 biological replicates, mean ± SD, *, p<0.05 (one-way ANOVA). **H.** LDH activity in the supernatant upon treatment of J774A.1 macrophages for two hours. n=3 biological replicates, mean ± SD, *, p<0.05 (one-way ANOVA). **I.** Normalized mRNA expression levels of IL10. n=4 biological replicates, mean ± SD, *, p<0.05 (one-way ANOVA). **J.** Representative Western blot cleaved caspase-3, arginase-1 and active p20 caspase-1 of J774A.1 macrophages untreated or incubated for four hours with different oils. **K.** Quantification of Western blots. n=3 biological replicates, mean ± SD, *, p<0.05 (one-way ANOVA).

To further analyze cell death in macrophages, J774A.1 cells were treated with different oils. All three vegetal oils increased cell death. In this case, the increase in Annexin V^+^, PI^+^ double positive cells was present for olive, corn and peanut oil. Moreover, necrotic cell death (Annexin V^−^, PI^+^) was also increased in the presence of peanut and corn oil (**Figure 6D**).

In order to further clarify whether macrophages die by apoptosis or necrosis we incubated J774A.1 cells with different oils to determine apoptotic and necrotic cell death. There was an increase in apoptotic cells in the case of incubation with olive oil and a mild increase in the presence of corn oil. 7-AAD incorporation (necrosis) was increased upon treatment with peanut and to a lesser extent upon treatment with corn and mineral oil (**Figure 6E-G**).

Another way to detect necrotic cell death is measuring lactate dehydrogenase (LDH) activity in the cell culture supernatant. Membrane disruption of necrotic cells allows release of cytosolic LDH. Peanut oil caused pronounced release of LDH. There was also LDH release from macrophages treated with olive and corn oil, however to a lesser degree (**Figure 6H**). Interestingly, the LDH release upon treatment with peanut oil could be strongly decreased by supplementing peanut oil with the polyunsaturated oleic acid (**Supplemental Figure 4A**).

We corroborated the different mechanisms of cell death further and observed that only treatment with olive oil induces cleavage of caspase-3 in macrophages, a major effector of apoptosis (**Figure 6J**). On the other hand, peanut oil-treated macrophages showed an increase in active p20 caspase-1, which is a marker for pyroptosis (**Figure 6J-K**), which has similar features as necrosis but is driven by caspase-1 activation (28). In contrast, olive oil-treated macrophages showed increased levels of IL10 and Arginase-1 indicating an anti-inflammatory switch towards resolution of inflammation (**Figure 6I-K**).

These results suggest that macrophages in contact with olive oil die by apoptosis, which facilitates the subsequent resolution of the inflammation, whereas peanut oil induces macrophage pyroptosis, which impairs the resolution of inflammation.

### *I.p.* injection of peanut oil impairs the resolution of inflammation in a peritonitis model

The results presented indicate that *i.p.* oil injection leads to a dramatic change in the peritoneal immune cell composition. Chronic inflammation is induced by peanut oil and this would potentially alter the outcome of experiments executed subsequently. One such example could be a peritonitis experiment in transgenic mice that had been injected *i.p.* before with tamoxifen in peanut oil to induce gene recombination. We decided to test this in experimental peritonitis model, in which mice received peanut oil by *i.p.* injection or by oral gavage as control. Three weeks later, thioglycolate was applied to mimic bacterial peritonitis (**Figure 7A**).

**Figure 7.**
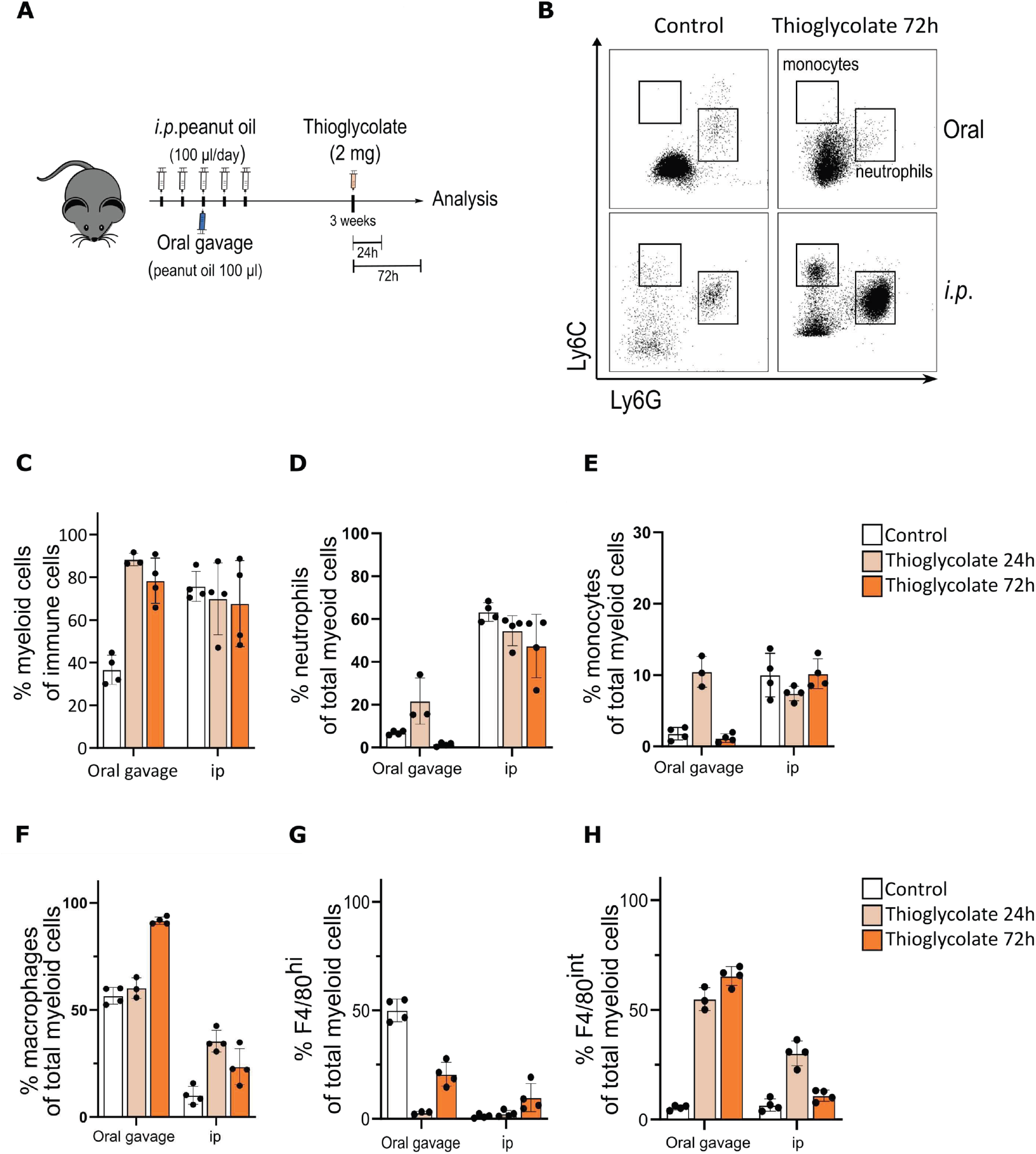
Intraperitoneal injection of peanut oil impairs the resolution of inflammation in a thioglycolate-induced peritonitis model. **A**. Schematic illustration of *i.p.* or oral oil administration followed by thioglicolate *i.p*. injection three weeks after treatment with oil. **B.** Representative blots of flow cytometry analysis of monocytes, neutrophils. **C**. Percentage of myeloid cells (CD45^+^CD11b^+^) in peritoneal lavage. **D**. Percentage of neutrophils (CD45^+^CD11b^+^Ly6G^+^Ly6C^int^) in peritoneal lavage. **E**. Percentage of monocytes in peritoneal lavage. **F**. Percentage of macrophages (CD45^+^CD11b^+^F4/80^+^) in peritoneal lavage. **G.** Percentage of macrophages CD45^+^CD11b^+^F4/80^hi^ in peritoneal lavage. **I.** Percentage of macrophages CD45^+^CD11b^+^F4/80^in^ in peritoneal lavage. n=3 mice for oral gavage followed by thioglycolate and analysis at 24 hours. All other groups n=4 mice.

It is known that thioglycolate initially induces a massive neutrophil and monocyte infiltration, followed by differentiation into macrophages that resolve inflammation by clearance of apoptotic cells (16). Consistently, mice that had received peanut oil by oral gavage had approximately 50% increase in myeloid cell numbers 24 and 72 hours after thioglycolate injection. However, mice pretreated *i.p.* with oil, already had high numbers of CD45^+^CD11b^+^ myeloid cells in peritoneal fluid at baseline and this was not further increased upon thioglycolate administration (**Figure 7C**). In mice treated by oral gavage, the number of monocytes and neutrophils in peritoneal fluid increased strongly upon thioglycolate injection and subsequently returned below baseline. This suggests that the first inflammatory response by these cells had already been cleared. However, in mice that had been *i.p.* injected with peanut oil there was a higher proportion of monocytes and neutrophils already under basal conditions, which was maintained after thioglycolate administration (**Figure 7B and D-E**).

Mice treated orally showed the expected increase in CD45^+^CD11b^+^F4/80^+^ macrophages 72 hours after thioglycolate injection. However, mice that had been injected *i.p.* with peanut oil had only few macrophages present in the peritoneum (approximately 12% of all myeloid cells) and these increased only marginally (**Figure 7F**). Orally treated mice showed disappearance of F4/80^hi^ macrophages 24 hours after thioglycolate administration and subsequent recovery, which was accompanied by an increase of F4/80^int^ macrophages (**Figure 7G-H**). This suggests that resident macrophages disappear after thioglycolate administration and get replaced by monocyte-derived macrophages as the inflammation resolves. Yet, in mice that received peanut oil *i.p*. there were only few F4/80^hi^ macrophages at baseline and there was only a minor increase in F4/80^int^ macrophages (**Figure 7G-H**). The lower presence of F4/80^int^ macrophages, together with the continuous influx of monocytes and neutrophils, suggests that resolution of inflammation cannot take place. As such, *i.p.* peanut oil injection leads to a dramatic change in the myeloid cell composition of the peritoneum that affects the outcome of subsequent experiments.

## Discussion

Animal experimentation requires careful planning and analysis to allow reproducibility and the possibility to translate basic research into successful clinical trials. Oil injection is frequently performed in animal research, in particular to deliver tamoxifen for inducible gene recombination (1, 2, 4). Oil is considered to be safe and non-toxic. However, few studies reported peritoneal inflammation after subcutaneous or intraperitoneal oil injection (6, 10, 29). To our knowledge, little is still known about changes in the immune cell composition and related reactions within the peritoneum upon intraperitoneal oil delivery. This work demonstrates that intraperitoneal injection of four different oils causes inflammation, foam cell formation and depletion of resident macrophages. However, the severity of inflammation strongly depends on the type of oil.

Within the peritoneum, the omentum plays a major role in recognition and encapsulation of pathogens (30). During this process, it expands, a feature which we observed after injection of the different oils. Interestingly, the applied oils were not completely resorbed even three weeks after injection into the peritoneal cavity. In particular larger amounts of peanut oil were still visible in the peritoneal fluid. The failed clearance can be assumed to prolong the phase of acute inflammation (17). Consistently, chronic xanthogranulomatous inflammation and fibrosis were observed, particularly upon peanut and mineral oil treatment. Myeloid cell infiltration and fibrosis were more severe in the omentum compared to the mesentery. This is consistent with the fact that the omentum is an immunological niche and the first organ to react against pathogens, but only when this inflammation becomes chronic it starts to affect the mesentery (20).

Mechanistically, the type of oil-induced macrophage cell death appears to determine whether inflammation gets resolved. For successful resolution of the inflammation, there is the need of efferocytosis, where macrophages engulf apoptotic cells (31). Non-resolving inflammation contributes substantially to the progression of atherosclerotic plaques and other chronic inflammatory diseases (18, 32, 33). Our data indicate that macrophages in contact with olive oil die by apoptosis, which facilitates efferocytosis-mediated resolution of inflammation. Conversely, peanut oil induces pyroptosis of macrophages. Excessive pyroptosis impairs the resolution of inflammation (34, 35). As such, peanut oil injection results in chronic peritoneal inflammation, whereas olive oil induces macrophage apoptosis, followed by efferocytosis and initiation of the resolution phase.

At the cellular level, all three vegetal oils were taken up by macrophages and caused foam cell formation. This change in the expression pattern has been observed in peritoneal macrophages isolated from obese mice, and blood monocytes from patients suffering from severe atherosclerosis (36–38). In principle, one could even use intraperitoneal peanut oil injection as a fast model to obtain viable foam cells for *in vitro* experiments. Future research will determine the potential of this model.

In conclusion, our study shows that intraperitoneal injection of different oils causes peritoneal inflammation and depletion of resident peritoneal macrophages. Whereas olive oil triggers macrophage apoptosis and resolution of inflammation, peanut oil induces pyroptosis and chronic non-resolved inflammation. This has important consequences for animal experiments. In a proof-of-principle approach, we demonstrated this in a thioglycolate-induced peritonitis model after peanut oil injection. To overcome such limitations, it is advisable to deliver lipophilic substances like tamoxifen by oral gavage instead of intraperitoneal injection.

## Acknowledgements

This work was funded by the Deutsche Forschungsgemeinschaft (DFG) project number 394046768 - SFB1366 projects C4 and Z2 (to A.F., C.M.), DFG project number 419966437 (to J.R.V.) and the Helmholtz Association (to A.F.).

