## Supplementary Figure for "Intraperitoneal oil application causes local inflammation with depletion of resident peritoneal macrophages" (from another version of this preprint: version 1, 2020-07-16)

### Supplementary figure legends

**Supplementary Figure 1. H&E staining of spleen after intraperitoneal injection of oil.** Representative confocal images showing H&E staining of spleen sections three weeks after oil injection or oral gavage. Spleens from untreated mice served as control. Scale bar, 200 μm.

**Supplementary Figure 2. Oil injection decreases the number of lymphocytes in peritoneal lavage. A**. Gating strategy of flow cytometry. **B**. Total cell number of B lymphocytes ($CD45^{+}CD19^{+}CD3^{-}$), T lymphocytes ($CD45^{+}CD19^{-}CD3^{+}$) and myeloid cells ($CD45^{+}CD19^{-}CD11b^{+}$) in peritoneal lavage. n=3 mice, mean ± SD, *, $p<0.05$ (one-way ANOVA).

**Supplementary Figure 3. Injection of saline does not generate inflammation in the peritoneum. A**. Schematic illustration of saline (0.9% NaCl) *i.p.* injection. Representative images at three weeks after treatment. **B**. Mouse peritoneum. **C**. Omentum. **D**. Mesentery. **E**. Total cell number in peritoneal lavage (PL) three weeks after injection. **F**. Gating strategy of flow cytometry. **G.** Flow cytometry analysis of total cell number of myeloid cells ($CD45^{+}CD11b^{+}$), neutrophils ($CD45^{+}CD11b^{+}Ly6G^{+}Ly6C^{-}$), monocytes ($CD45^{+}CD11b^{+}Ly6G^{-}Ly6C^{+}$) and macrophages ($CD45^{+}CD11b^{+}F4/80^{+}$). All data represent n=5 mice. Bar graphs show mean ± SD.

**Supplementary Figure 4. Oleic acid supplementation prevents peanut oil-induced cytotoxicity.** LDH activity in the supernatant of J774A.1 macrophages upon treatment for two hours with 100 μl DMEM (untreated), 5 μl of peanut oil in 100 μl DMEM (Peanut) or 5 μl of peanut oil and 200 μM of oleic acid in 100 μl DMEM (Peanut + Oleic Acid). Data represent n=3 biological replicates. Bar graph shows mean ± SD, *, $p<0.05$ (one way ANOVA).

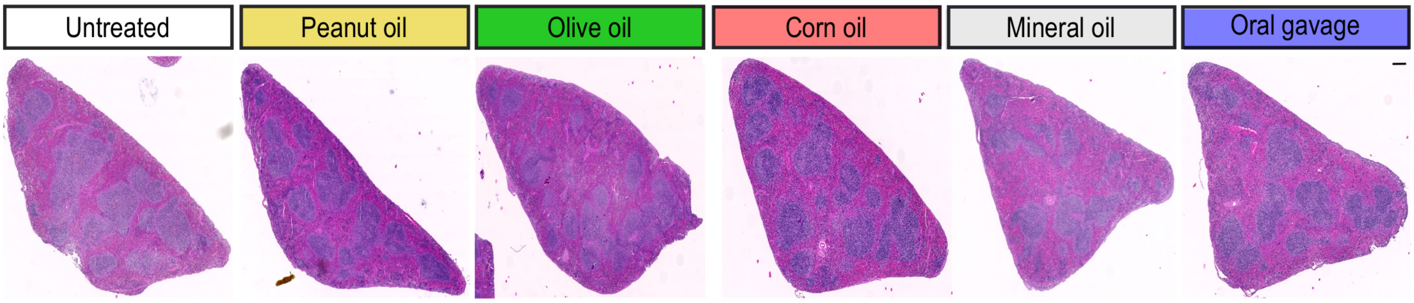


**Supplementary Figure 1**.

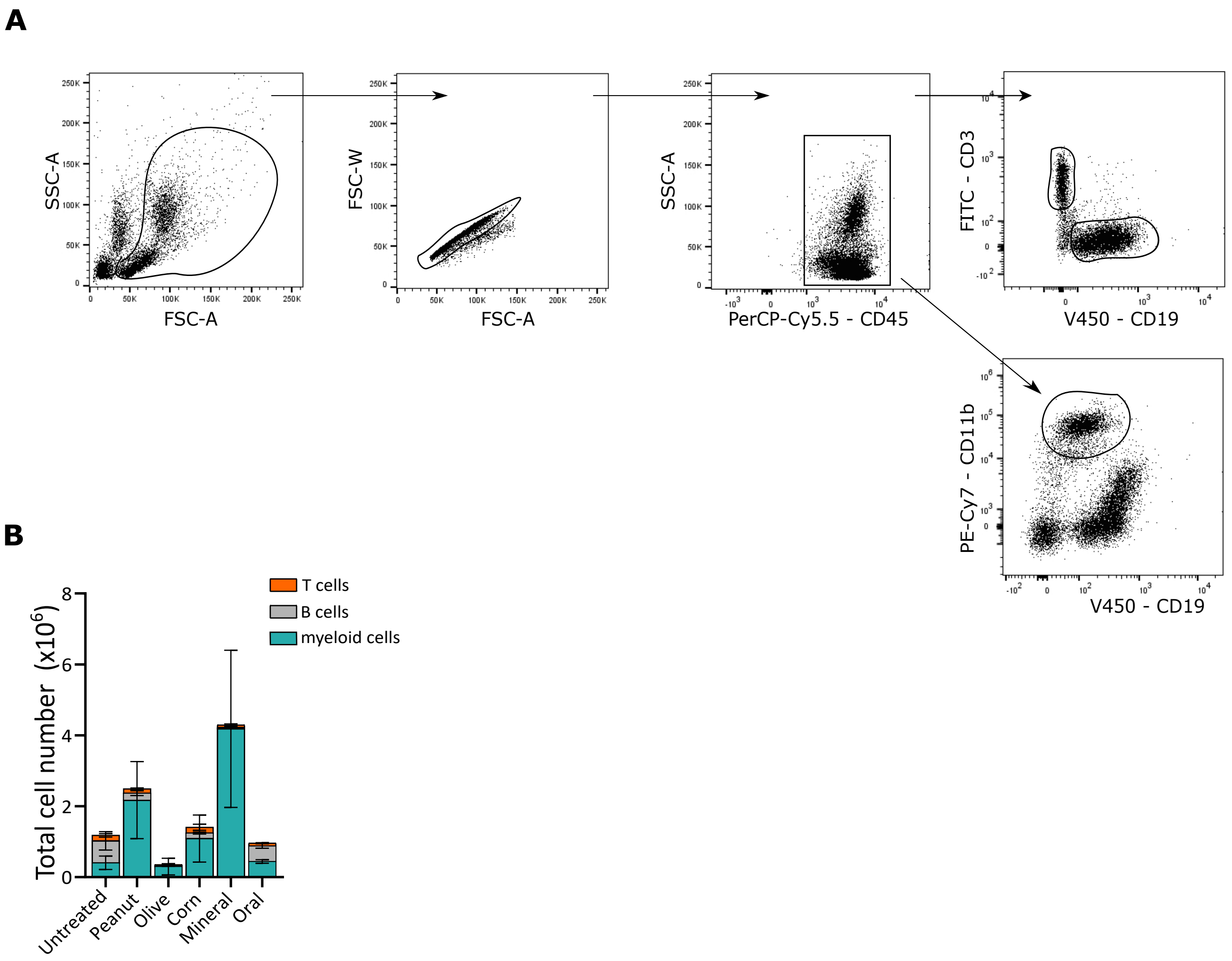


**Supplementary Figure 2**.

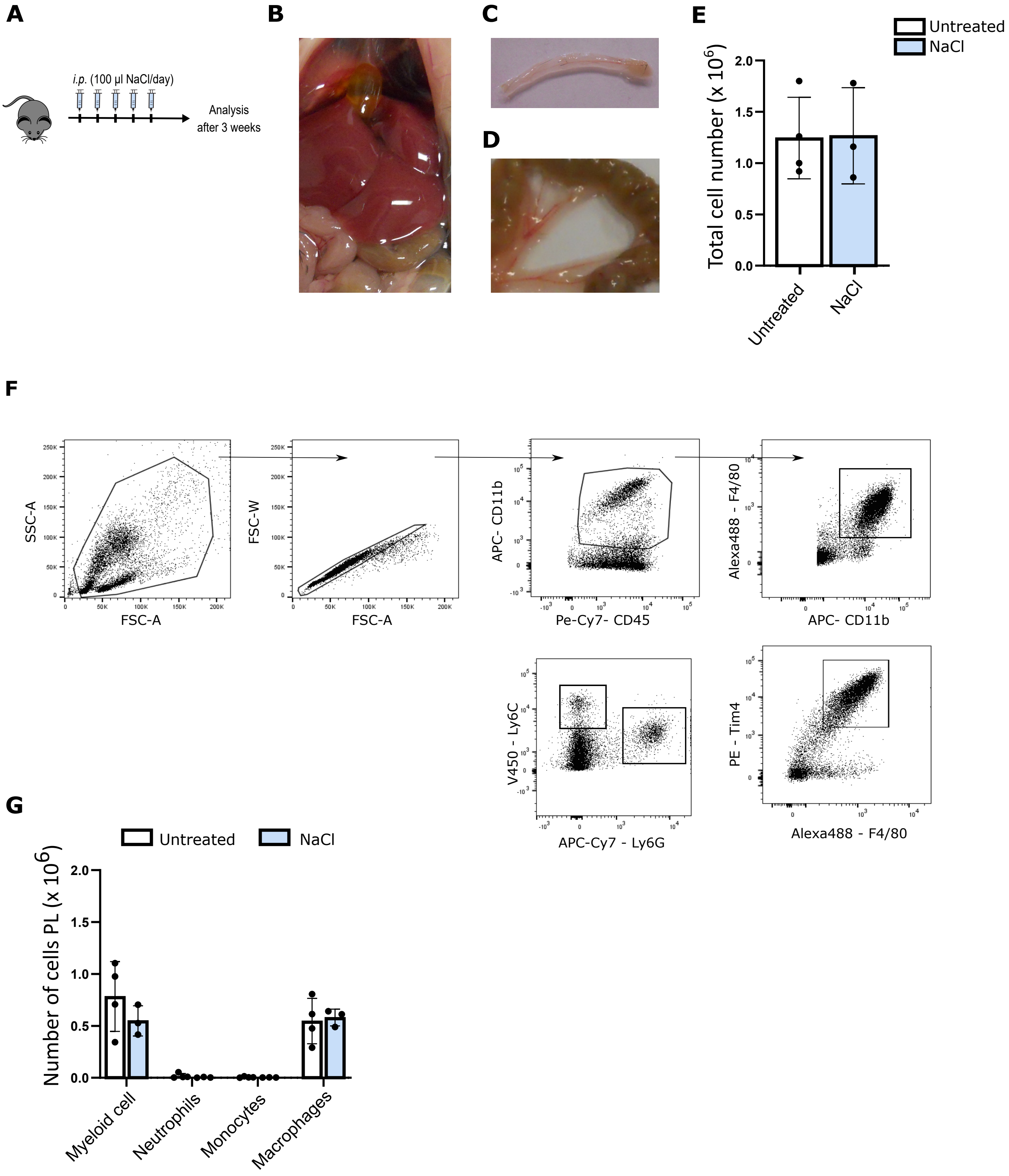


**Supplementary Figure 3**

**A**

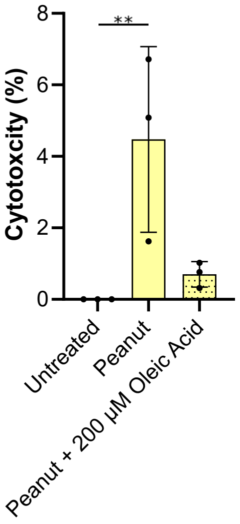


**Supplementary Figure 4**.
